# CoNekT: an open-source framework for comparative genomic and transcriptomic network analyses

**DOI:** 10.1101/255075

**Authors:** Sebastian Proost, Marek Mutwil

## Abstract

The recent accumulation of gene expression data in the form of RNA sequencing creates unprecedented opportunities to study gene regulation and function. Furthermore, comparative analysis of the expression data from multiple species can elucidate which functional gene modules are conserved across species, allowing the study of the evolution of these modules. However, performing such comparative analyses on raw data is not feasible for many biologists. Here, we present CoNekT (Co-expression Network Toolkit), an open source, user-friendly web server, that contains user-friendly tools and interactive visualizations for comparative analyses of gene expression data and co-expression networks. These tools allow analysis and cross-species comparison of (i) gene expression profiles; (ii) co-expression networks; (iii) co-expressed clusters involved in specific biological processes; (iv) tissue-specific gene expression; and (v) expression profiles of gene families. To demonstrate these features, we constructed CoNekT-Plants for green alga, seed plants and flowering plants (*Picea abies, Chlamydomonas reinhardtii*, *Vitis vinifera*, *Arabidopsis thaliana*, *Oryza sativa*, *Zea mays* and *Solanum lycopersicum*) and thus provide a web-tool with the broadest available collection of plant phyla. CoNekT-Plants is freely available from http://conekt.plant.tools, while the CoNekT source code and documentation can be found at https://github.molgen.mpg.de/proost/CoNekT/.

## Introduction

With the continuous improvement of sequencing technologies, the cost to generate a genome sequence has decreased nearly 8000-fold during the last decade (https://www.genome.gov/sequencingcostsdata/). Due to these improvements, RNA sequencing (RNA-Seq) became the method of choice to study transcript abundance. RNA-Seq allows detection of differentially expressed genes (1), assembly of coding sequences *de novo* in the absence of a reference genome (2), construction and analysis of expression atlases (3–5) and co-expression networks which can guide gene-function predictions (6, 7). Combined with comparative genomics, these approaches can also be used to study transcriptional differences to understand phenotypic variation within and between species (8–10).

While various tools exist to browse expression profiles and co-expression networks (8, 11–14), they are often limited to few species and closed-source, which prevents users to create custom versions including their own data. To this end, we developed CoNekT (Co-expression Network Toolkit, https://github.molgen.mpg.de/proost/CoNekT). As CoNekT is open-source and available under the MIT license, researchers can create new online or in-house instances for their own data and expand CoNekT with features relevant to their research. To demonstrate the usefulness of the platform, we present CoNekT-Plants (http://conekt.plant.tools), which allows comparative analyses of six land plants and alga.

## Materials and Methods

### Implementation and Interface

CoNekT consists of two components; (i) a python-flask backend which processes requests, fetches data from the database, provides search functionality and serves web pages and (ii) a front-end which includes various interactive visualizations based on charts.js, cytoscape.js (15), and phyD3.js (16). The Bootstrap CSS library is used to style all pages and is paired with jQuery.js and qtip2.js to add various dynamic elements such as tooltips and popups. Font-awesome is included for different glyphs and icons. A full overview of all included libraries in both the back- and front-end is included in the online documentation (https://github.molgen.mpg.de/proost/CoNekT/).

CoNekT contains pages for species, genes, gene families, co-expression clusters and neighborhoods, and others. These pages, in turn, contain graphs, tables, and links relevant to the page. For example, gene pages indicate the gene’s (i) description, (ii) gene family, (iii) phylogenetic tree, (iv) cDNA and protein sequences, (v) expression profile, (vi) co-expression neighborhood and cluster, (vii) similar neighborhoods in other species and (viii) Gene Ontology information (inferred by experimental evidence, InterProScan and co-expression network neighborhood)(Figure 1). A detailed description of all available features and instructions of how to deploy your own CoNekT web-server can be found at: https://github.molgen.mpg.de/proost/CoNekT/.

**Figure 1.**
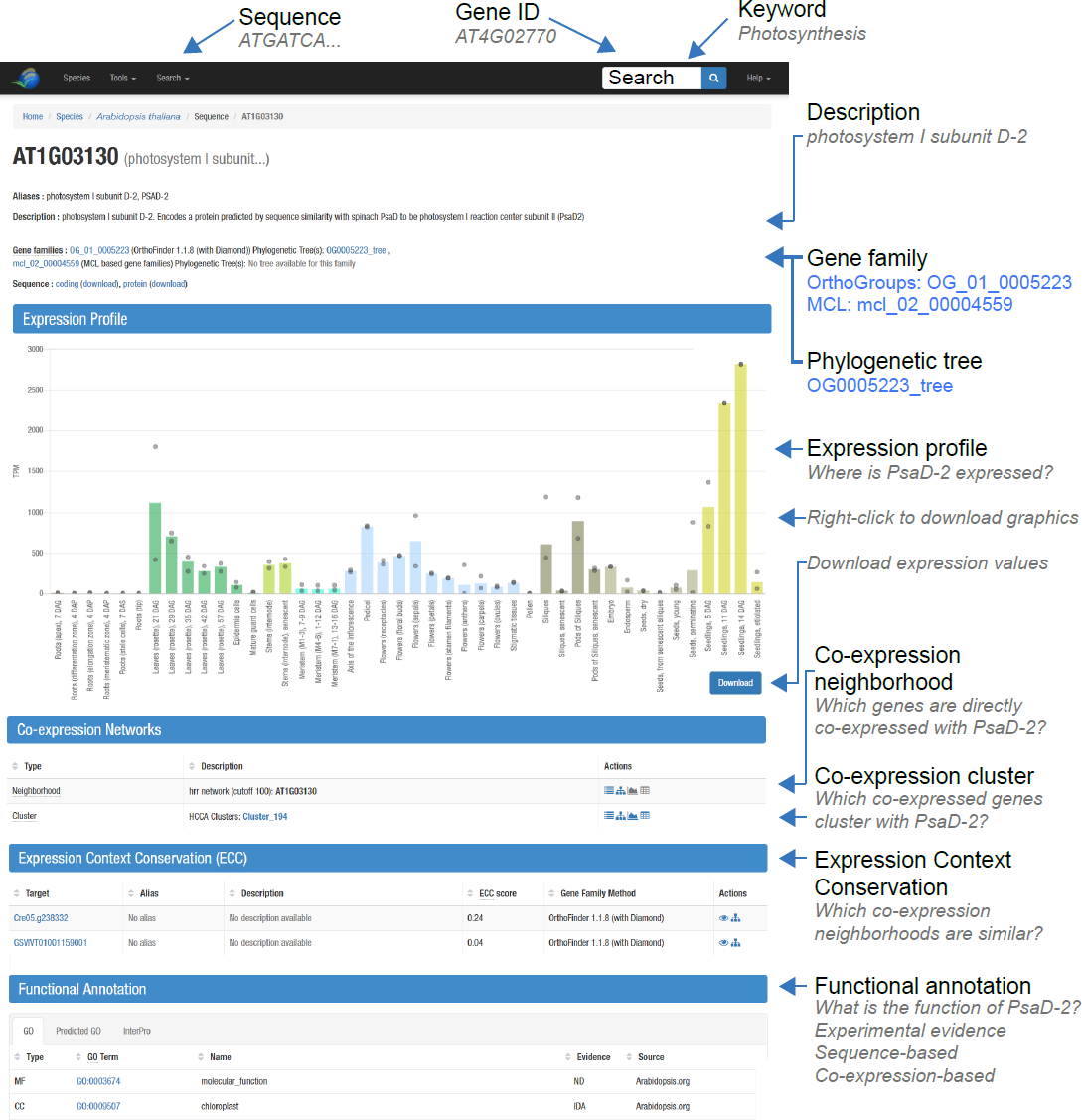
**Gene page contents exemplified with Arabidopsis *PsaD-2*.** The gene page provides information (as tables) and links (in blue) specific to the gene. The links allow quick access to the co-expression neighborhood, cluster, gene family and phylogenetic tree of *PsaD-2.*

### Data Acquisition for CoNekT-Plants

To demonstrate the web-server, we introduce CoNekT-Plants, which contains data from seven species (Table 1), including green alga *Chlamydomonas reinhardtii*, gymnosperm *Picea abies*, two monocots (*Oryza sativa*, *Zea mays*) and three dicotyledonous plants (*Vitis vinifera*, *Arabidopsis thaliana*, and *Solanum lycopersicum*). For each species, publically available RNA-Seq data was obtained through the Sequence Read Archive’s ‘Run Selector’ (https://www.ncbi.nlm.nih.gov/sra/)(17). These samples were downloaded, converted to fastq files (using SRATools, https://www.ncbi.nlm.nih.gov/books/NBK158900/) and processed using LSTrAP (6), which maps reads to the genome using TopHat (18) and determines transcript abundance for each gene using HTSeq-count (19). The mapping statistics included in LSTrAP were used to detect and discard samples that showed either (i) low mapping to the genome (< 65%), (ii) low mapping to coding sequences (< 40%) or (iii) too few useful reads (less than 8M reads mapping to the genome). Additionally, using LSTrAP’s heatmap tool, the output was screened for outliers, which were removed from the final dataset. The remaining samples were used to construct expression matrices and co-expression networks. For *Arabidopsis thaliana*, experimentally determined functional annotation (Gene Ontology terms) was obtained from www.arabidopsis.org. Additionally, for all species, InterProScan v5.18 (20) was used to detect protein domains and obtain predicted functional annotation. To obtain orthologs, OrthoFinder v1.1.8 (21) was used to group genes into orthogroups and construct phylogenetic trees, using Diamond to determine sequence similarities with settings at default values (22). Sequence similarities reported by Diamond were clustered using MCL to group homologous genes into gene families (23). Note that all abovementioned steps can be performed in LSTrAP, and the output can be directly used in CoNekT.

**Table 1.**
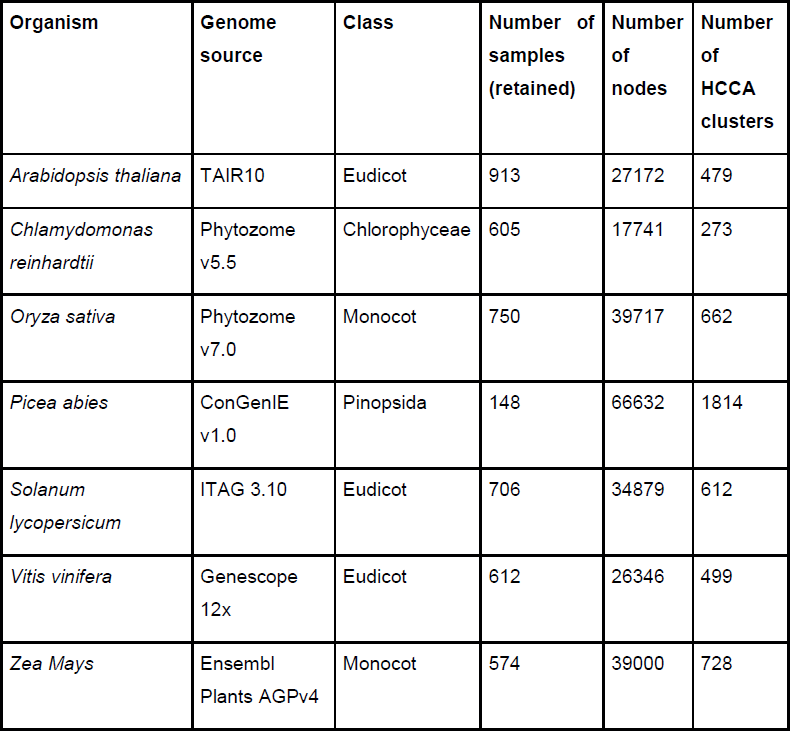
**Species included in CoNekT-Plants.** The table indicates the genome source, phylogenetic class, number of RNA-seq samples that passed the LSTrAP quality control, number of nodes (genes) and the number of co-expression clusters identified by HCCA algorithm.

Using CoNekT’s graphical admin interface, the expression and genomic data were added to the platform (see instructions on https://github.molgen.mpg.de/proost/CoNekT/). Through the same interface, multiple analyses were started, such as (i) the Heuristic Cluster Chiseling Algorithm (HCCA), to find clusters of co-expressed genes in the networks (24); (ii) Gene Ontology term over-representation to elucidate the functional annotation of co-expression clusters (reported as enrichment fold-changes and p-values); (iii) identification of similar co-expression network clusters within and across species, and others.

## Results and discussion

### Querying CoNekT

CoNekT features three modes to search for relevant content. First, the keyword search, available from the landing page and upper right corner (Figure 1), accepts gene IDs (e.g. *At4g32410*), Gene Ontology term IDs (e.g. GO:0008810), keywords (e.g. “cellulose”) and InterPro domains (e.g. cellulose_synt), and returns genes, GO terms and InterPro domains that match the query. Second, the advanced search function available in Search/Search(advanced) menu on the top of the page can be used to retrieve genes with a specific combination of functional annotation, GO IDs and/or InterPro domains. Third, relevant genes can be retrieved by sequence similarity with BLAST (Search/BLAST).

### Gene Expression Profiles

The pattern of gene expression can reveal where and when a specific gene is active and thus can suggest the gene’s function. For example, uncharacterized genes with specific expression in roots might be essential for root development. To visualize gene expression, expression levels, defined here as Transcripts Per Kilobase Million (TPM), were grouped by tissue, condition and/or developmental stage for each gene (Figure 1). These profiles can be exported as png/jpg graphics or as a table.

CoNekT allows comparisons of gene expression across species, where average expression in predefined organs is shown. CoNekT-Plants was configured to show gene expression in roots/rhizoids, leaves, stems, female reproduction (containing ovaries, pistrils), male reproduction (containing pollen, anthers(and flower/seeds/spores. This cross-species comparative expression analysis is available from Tools/Heatmap/Comparative window or by clicking on “View comparative expression as a heatmap:” link on a gene family page. We illustrate such a heatmap with photosystem I subunit family D (PsaD, http://conekt.plant.tools/family/view/5224), which is involved in photosynthesis (25). The heatmap can be accessed by clicking on the “row-normalized” link in “View comparative expression as heatmap:” line. As expected of photosynthesis-related genes, they show the highest expression in leaves and virtually no expression in roots or male reproduction, which contains non-photosynthesizing pollen and anthers (Figure 2A). While this example illustrates that the PsaD family genes have conserved expression, the heatmap could be used to rapidly identify genes with changed expression.

**Figure 2.**
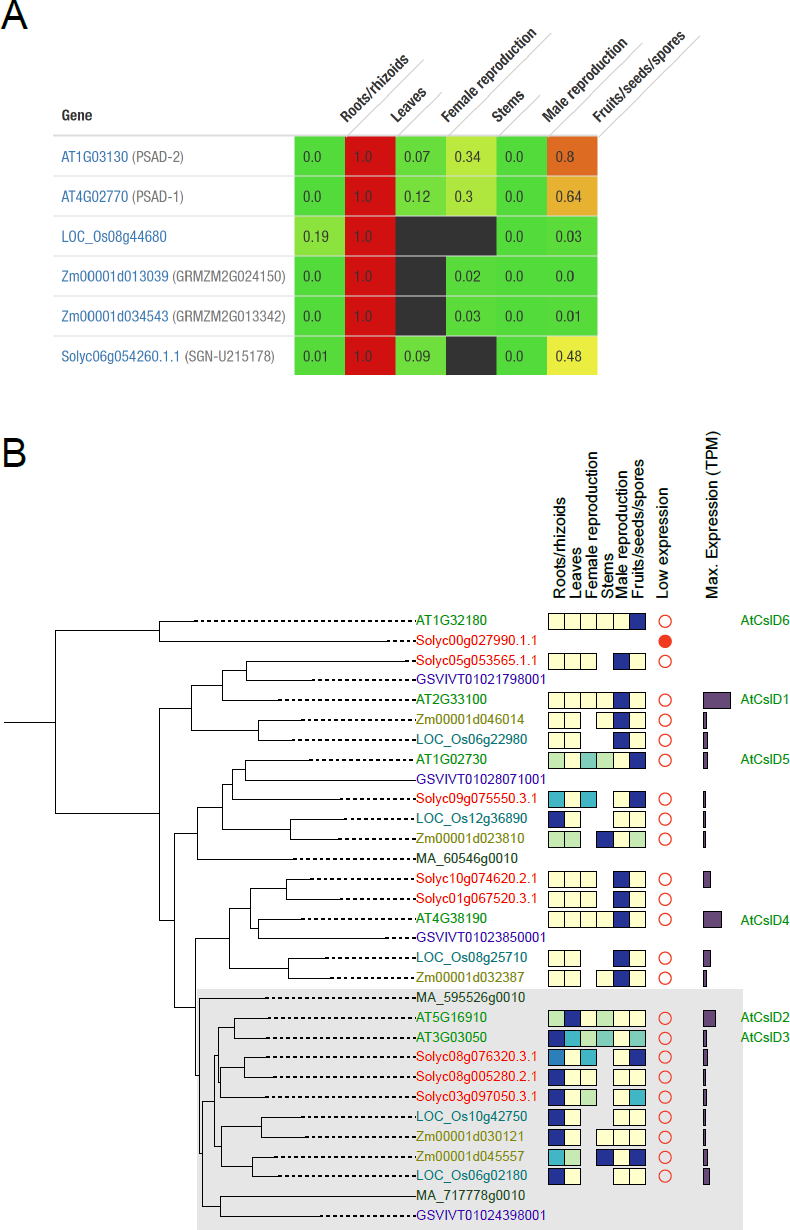
**Comparative expression analysis.** A) Expression profile of Arabidopsis (genes starting with AT), rice (LOC), maize (Zm) and tomato (Solyc) PsaD gene family in roots, leaves, female reproduction (e.g. ovaries, stigma), stems, male reproduction (e.g. pollen, anthers) and fruits. The expression values of each gene were normalized by diving by their maximum, and range from 1 (red, maximum expression) to 0 (green, no expression). Missing expression data is shown with a black box (e.g. female reproduction and stems for rice). B) Phylogenetic tree of the Cellulose Synthase-like D (CSLD) gene family. The heatmap shows the expression level in different tissues, full red dots show genes with low-expression and the bar on the right indicates the maximum expression level (TPM). The color of a gene identifier indicates the species. The added gray box contains genes that have shifted towards being expressed in roots. Note that OrthoFinder tree nodes do not contain bootstrap values, and should be interpreted with care. Missing data is indicated by absent box; for example, spruce has insufficient expression data to provide an informative expression.

### Expression specificity

Since gene expression patterns can reveal gene function, extracting genes expressed specifically in a given organ, tissue or condition can be used to predict gene function. To detect expression specificity, CoNekT uses specificity measure (SPM, ranges between zero and one, where one indicates the gene is exclusively expressed in the tissue)(26), Tau (high values indicate that a profile is specific in a tissue), and entropy (indicates how much a profile fluctuates across all tissues, where genes with very specific or very stable expression have low entropy)(27). The application of one or more of these metrics allows users to search for expression profiles specific for one tissue or condition (27). To illustrate the tool, we selected “Tools/Find specific profiles,” chose “Tissue specificity” as method and selected Arabidopsis “Meristems” as condition. The output is returned as a table, where rows are genes, columns contain descriptions, and the SPM, entropy and Tau values. For meristems in Arabidopsis, the tool returned a table with 146 genes with SPM>0.85 (default value) and showed known factors influencing meristem and flower development, such as *LEAFY*, *CLAVATA3*, *AINTEGUMENTA*, *DORNROSCHEN*, and others (28–31)(See Supplementary Table 1 for full list). Thus, the presence of these known factors indicates that the tool can retrieve relevant genes.

### Comparative expression specificity

When orthologous genes are specifically expressed in similar tissues/organs across different species, it further strengthes the evidence of their importance for the function of that tissue, as conserved expression profiles are less likely to appear by chance (8, 9, 32). Furthermore, orthologs that show conserved expression are more likely to be functionally equivalent, which can be used to resolve often unclear phylogenetic relationships caused by gene duplications (8, 9). To this end, the “Tools/Compare specificities” tool compares two lists of specifically expressed genes within or across species. As an example, we compared orthologs specifically expressed in Arabidopsis and rice pollen (Selected method “Tissue specificity”, default SPM), which revealed 103 orthogroups that were specifically expressed in pollen of the two plants. The results include a set of well-known genes related to pollen development and fertilization, such as *CSLD1*, *CSLD4*, *COBL10*, *LIP1*, *LIP2*, *TIP5;1*, *FH3*, *AKT6* (33–38), but also a host of other genes potentially important for these processes in both species (See Supplementary Table 2 for full list). Thus, similarly to the tool above, this feature can reveal genes relevant for the development of a tissue of interest, with the additional advantage of highlighting conserved and biologically relevant genes.

### Phylogenetic and expression analysis

Phylogenetic trees provide the most detailed view of speciations and duplications, and their timing, between homologous genes. However, phylogenetic trees not reveal sub- or neo-functionalization of genes that might be apparent when investigating expression data (10, 39, 40). To remedy this, CoNekT combines interactive phylogenetic trees with the comparative expression profile heatmaps, which allows elucidating potential sub- or neo-functionalization events.

To demonstrate the usefulness of the tool, we show a tree of Cellulose Synthase-like D (CslD) gene family, involved in tip growth in plants (Figure 3)(33). To obtain the tree, we first navigated to the page of *AtCslD1* (*At2g33100,* http://conekt.plant.tools/sequence/view/45080), which is involved in pollen tube growth (33), and clicked on a link to the family’s phylogenetic tree (OG0000579_tree, http://conekt.plant.tools/tree/view/12121). The tree revealed that most of the CslD family genes are either expressed in roots or male reproductive tissues (Figure 2B). For example, *AtCslD3* (*At3g03050*) shows high expression in roots, which is in line with the gene’s involvement in root hair growth (33). Furthermore, genes from other plants that are found in the *AtCslD3* clade (indicated by the gray box) show root-specific expression, suggesting that these genes are also involved in root hair growth. Conversely, most genes in the *AtCslD1* (*At2g33100*) and *AtCslD4* (*At4g38190*) clade show predominantly male reproduction-specific expression, which is in line with their function in pollen tube growth (33). *AtCslD5* (*At1g02730*) was shown to be involved in cell plate formation (41), suggesting that the clade the gene is found in has gained new function (Figure 2B). Finally, since the oldest lineage in the root hair and pollen tube clades is spruce (genes starting with *MA_*), we can postulate that the gene duplication that created the sub-specialized pollen tube / root hair genes took place in the common ancestor of seed plants. To conclude, the combination of phylogenetic trees and expression can be used to identify functional innovations found in gene families.

**Figure 3.**
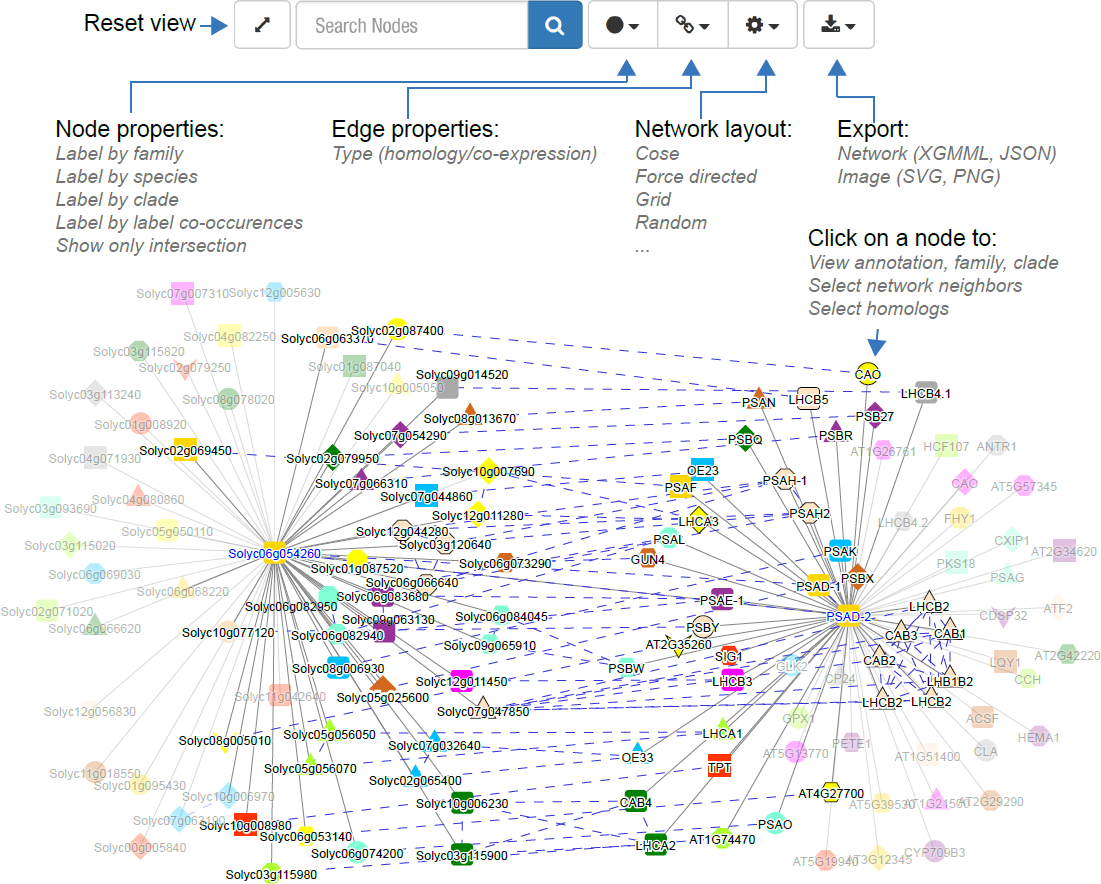
**Comparative network analysis of *AtPsaD-2* and Solyc06g054260.** *AtPsaD-2* and tomato ortholog *Solyc06g054260* are shown together with their co-expression neighborhood (co-expressed genes are connected using solid gray lines). Nodes with the same shape and color are members of the same OrthoGroup. Orthologs found in both neighborhoods are connected with dashed blue lines. The indicated menus are used to change the node and edge labels, network layout and export the networks as images and Cytoscape-compatible data. For clarity, non-conserved nodes were made semi-transparent.

### Comparative co-expression network analyses

Genes with similar expression profiles (co-expressed genes) are often functionally related, and consequently, co-expression analysis is a robust method for gene function prediction (8, 12, 42, 43). Co-expressed genes can be represented as a network, where nodes represent genes and edges are drawn between co-expressed nodes. Co-expression networks in CoNekT are based on Highest Reciprocal Rank (HRR) metric score of 100 or better (8), which is related to a robust rank-based metric used to identify co-expressed genes (44). Groups of densely connected genes, called co-expression clusters, were detected using the Heuristic Cluster Chiseling Algorithm (24). Co-expression patterns can be conserved across species (even over large phylogenetic distance)(45–47), and this property can be used to transfer functional knowledge from one species to another (8, 48–50). Recently, it was demonstrated that subsections of the co-expression network comprised of functionally related genes, also known as gene modules, were duplicated within one species (40). To detect these conserved and duplicated modules, CoNekT uses the Expression Context Conservation (ECC) value (9) to detect which network regions contain more gene families than expected by chance (51).

Users can visualize a gene and its direct co-expression partners (neighborhood) or all genes within a co-expression cluster (Figure 3). The interactive networks provide an intuitive interface that allows nodes to be colored based on various parameters (e.g., gene family, phylogenetic clade and others) and can be searched by gene name, alias or annotation. Edges can be colored and shown/hidden by edge weight. Furthermore, different graph layouts are supported, and the networks can be exported as vector (SVG) or pixel-based (PNG) graphics.

To exemplify a comparative co-expression analysis, we used Arabidopsis *PsaD-2*, which is part of photosystem I complex. On the *PsaD-2* gene page (*At1g03130*, http://conekt.mpimp-golm.mpg.de/pub/sequence/view/35615), a tomato ortholog (*Solyc06g054260.1.1*) with an ECC score of 0.33 was found. By clicking on “View ECC pair as graph” glyph, CoNekT detected conserved photosynthetic components of photosystem I (gene IDs starting with PSA), II (gene IDs starting with PSB), Light Harvesting Complex (gene IDs starting with LHC) in both genes’ neighborhoods (orthologs are connected by dashed lines, Figure 3).

### Searching for functionally enriched clusters across species

To detect co-expression clusters containing genes associated with specific functions, CoNekT precalculates GO enrichment for all clusters, which can be searched using the ‘Tools\Find enriched clusters’ feature. For instance, looking for clusters enriched for GO term ‘reproduction’ in all seven species found in CoNekT-Plants yielded 18 clusters significantly enriched for this term. By clicking on “Compare profiles in this cluster” icon in “Action” menu, users can quickly screen which cluster is acting in the tissues of interest. Such search revealed that *Arabidopsis thaliana* cluster 17 and maize cluster 6 were significantly enriched for genes involved in reproduction (adjusted P-value<0.01) and expressed in pollen and anthers (Supplementary Table 3), indicating that these clusters are involved in a male reproductive process. Such analysis is therefore a good starting point to identify genes relevant for a biological process of interest.

## Conclusions

CoNekT is a modern web-platform that provides an intuitive interface for combining large-scale expression data with functional and genomic information. This allows users to extract tissue-specific genes, to compare tissue-specific transcriptomes between species and to leverage co-expression networks to predict gene function. These networks can be compared in a broad phylogenetic context. As CoNekT is open-source, researchers can create a version which includes their own RNA-Seq data and disseminate this online. Expert users can dive in the code and implement advanced features designed to answer their specific research questions, without having to re-implement core components such as gene families, expression profiles and co-expression network browsers.

## Data availability

The co-expression networks are available for download at http://conekt.plant.tools/species/. Source code and documentation can be found on https://github.molgen.mpg.de/proost/CoNekT/

## Supplementary data

Supplementary Data are available at NAR online.

## Acknowledgements

CoNekT-Plants is hosted at the Max Planck Institute of Molecular Plant Physiology MPIMP in Potsdam-Golm, Germany and we would like to thank Andreas Donath for tech support. Furthermore, we would like to thank EvoRepro members for testing and providing valuable feedback, especially Jörg Becker, Ann-Catherin Lindner, David Twell, and Mark Johnson, and Camilla Ferrari for proofreading the manuscript. Finally, we would like to express our gratitude to Maximilian Funk for help with licenses.

## Author contributions

CoNekT was designed and implemented by S.P who also prepared the data and build CoNekT-Plants with input from M.M. Both S.P and M.M. wrote the manuscript.

